# Human Foveal Cone Photoreceptor Topography and its Dependence on Eye Length

**DOI:** 10.1101/589135

**Authors:** Yiyi Wang, Nicolas Bensaid, Pavan Tiruveedhula, Jianqiang Ma, Sowmya Ravikumar, Austin Roorda

## Abstract

We provide the first measures of foveal cone density as a function of axial length in living eyes and discuss the physical and visual implications of our findings. We used a new generation Adaptive Optics Scanning Laser Ophthalmoscope to image cones at and near the fovea in 28 eyes of 16 subjects. Cone density and other metrics were computed in units of visual angle and linear retinal units. The foveal cone mosaic in longer eyes is expanded at the fovea, but not in proportion to eye length. Despite retinal stretching (decrease in cones/mm^2^), myopes generally have a higher angular sampling density (increase in cones/deg^2^) in and around the fovea compared to emmetropes, offering the potential for better visual acuity. Reports of deficits in best-corrected foveal vision in myopes compared to emmetropes cannot be explained by increased spacing between photoreceptors caused by retinal stretching during myopic progression.

## Introduction

There has been a rapid increase in prevalence of myopia, of all magnitudes, in the period between 1971-1972 and 1999-2004 (Vitale, Sperduto, & Ferris, 2009). Across sub-populations grouped by race, ethnicity and gender, several studies report axial length of the eye to be the primary variable related to myopia (Gonzalez Blanco, Sanz Ferńandez, & Muńoz Sanz, 2008; He et al., 2015; Iyamu, Iyamu, & Obiakor, 2011). Increased axial length is associated with retinal stretching and thinning of posterior segment layers and the choroid (Fujiwara, Imamura, Margolis, Slakter, & Spaide, 2009; Harb et al., 2015) and is associated with sight-threatening, often irreversible pathologies of the retina (Morgan, Ohno-Matsui, & Saw, 2012; Verkicharla, Ohno-Matsui, & Saw, 2015). Even without any detectable pathology, the structural changes associated with eye growth ought to have functional consequences for vision.

### What Do We Know About Functional Deficits in Myopia?

One might expect that eye growth would stretch the photoreceptor layer and would increase the spacing between cones, causing a longer eye to more coarsely sample an image relative to a shorter eye. However the situation is not that simple; the axial elongation associated with eye growth is accompanied by magnification of the retinal image (Strang, Winn, & Bradley, 1998). If the enlargement of the retinal image exactly matched the stretching of the cone mosaic, then eyes of different lengths would sample the visual field similarly. In fact, in large scale studies, myopes generally attain reasonably good visual acuity with optical correction (He et al., 2004; Jong et al., 2018).

However, more careful inspection reveals that myopes generally (6 out of 9 studies) have poorer angular resolution and have uniformly (3 out of 3 studies) poorer retinal resolution. Table 1 summarizes published results from psychophysical foveal tasks.

**Table 1:**
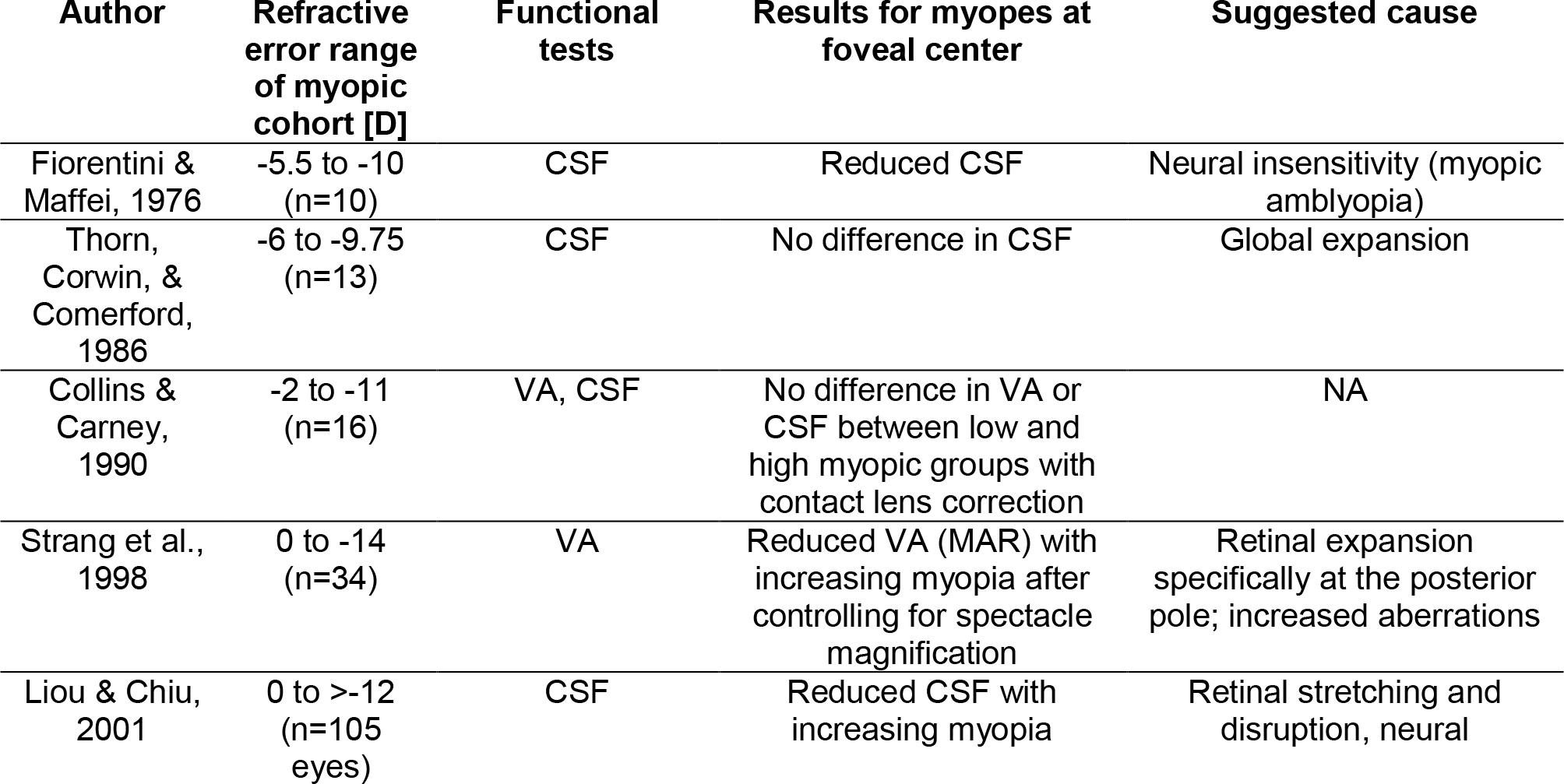
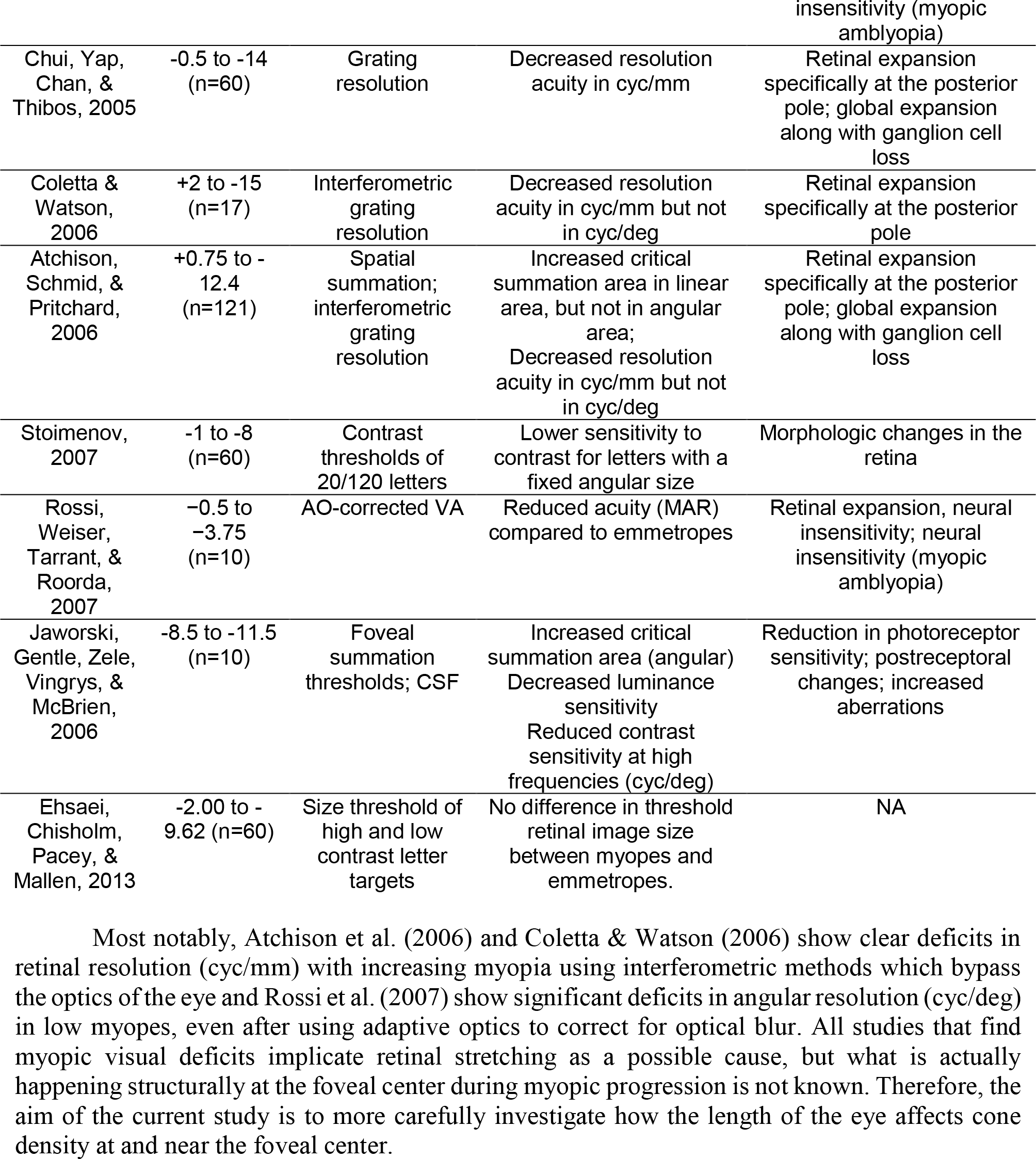
Summary of studies investigating foveal spatial vision and sensitivity tasks in myopia.

Most notably, Atchison et al. (2006) and Coletta & Watson (2006) show clear deficits in retinal resolution (cyc/mm) with increasing myopia using interferometric methods which bypass the optics of the eye and Rossi et al. (2007) show significant deficits in angular resolution (cyc/deg) in low myopes, even after using adaptive optics to correct for optical blur. All studies that find myopic visual deficits implicate retinal stretching as a possible cause, but what is actually happening structurally at the foveal center during myopic progression is not known. Therefore, the aim of the current study is to more carefully investigate how the length of the eye affects cone density at and near the foveal center.

### Models for How Photoreceptors Change with Eye Growth

Two types of cone densities will be discussed in this study. Linear density quantifies how many cones are within a fixed area, in square mm, and serves as a way to evaluate physical retinal stretching caused by eye growth. Angular density quantifies how many cones are within one degree visual angle, (the visual angle is measured from the secondary nodal point of the eye). Angular density serves as a way to evaluate the visual implications of eye growth as it governs the sampling resolution of the eye.

Figure 1 illustrates three models, along the lines of Strang et al. (1998), of how photoreceptor structure might be affected by myopic eye growth. In the first model, called the **global expansion model**, the retina is proportionally stretched with increasing axial length - cones are more spaced out in longer eyes - and linear density decreases with eye length. Assuming that the secondary nodal point remains at a fixed position relative to the anterior segment, the number of cones within a fixed angular area will remain constant. Therefore, angular cone density will be constant with eye length. In the second model, called the **equatorial stretching model**, the posterior retina simply moves axially further from the anterior segment of the eye so that the linear density does not change with eye length. Since the retina is moving further from the secondary nodal point, more cones will fall within a fixed angular area and the angular cone density will increase with eye length. The final model, called the **over-development model**, describes a structural photoreceptor change that mimics the changes that occur during development (Springer & Hendrickson, 2004) whereby the photoreceptors continue to migrate towards the fovea as the eye grows. In this scenario, longer eyes will show both increased linear cone density and an even steeper increase in angular cone density. The model is motivated by observations of increased linear cone density in the foveas of marmosets that underwent lens-induced eye growth (Troilo, 1998).

**Figure 1:**
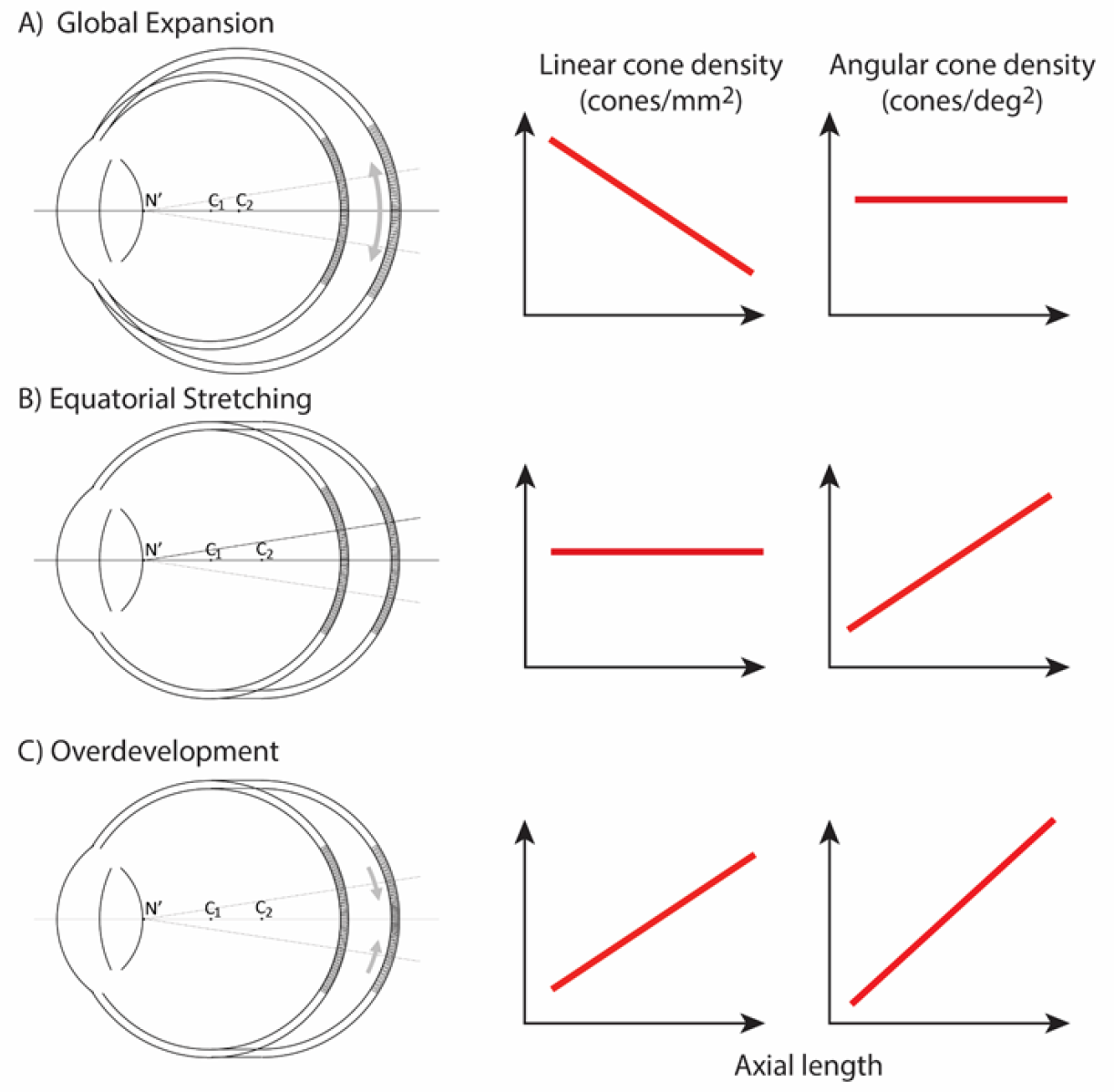
3 models of myopic eye growth: (A) Global expansion shows an eyeball that is proportionally stretched. (B) The equatorial stretching model indicates a growth model where the fovea stays rigid and unaffected as the eye grows. (C) The over-development model shows that myopic eye growth is similar with developmental eye growth where photoreceptors continue to migrate towards the fovea as the eye grows.

### Previous Studies of Cone Spacing with Axial Length

The most definitive studies of cone spacing as a function of axial length are done through direct imaging of the retina – wherein sharp images of the cones are enabled through the use of adaptive optics, a set of technologies that actively compensate the blur caused by aberrations of the eye (Liang, Williams, & Miller, 1997). Combined with confocal scanning laser ophthalmoscopy (Webb, Hughes, & Delori, 1987), adaptive optics offers the highest contrast *en face* images of the foveal photoreceptor mosaic ever recorded in vivo (Dubra et al., 2011; Roorda et al., 2002).

Despite continued advances in image quality, previous studies investigating cone packing and eye length have not made their measurements at the foveal center, the most important region for spatial vision but the most difficult to image owing to the small size of photoreceptors. There are a number of studies on cone packing and eye length (Chui, Song, & Burns, 2008; Elsner et al., 2017; Kitaguchi et al., 2007; Li, Tiruveedhula, & Roorda, 2010; Obata & Yanagi, 2014; Park, Chung, Greenstein, Tsang, & Chang, 2013) and here we summarize the published results that are most relevant to our study. Chui et al. (2008) investigated angular and linear cone density at 1 mm and 3 degrees eccentricity. They found a significant decrease (P<0.05) in linear cone density as a function of eye length at 1mm (which, by angular distance, is closer to the fovea in a longer eye than in a shorter eye) in all directions except in the nasal retina. They found that the angular cone density at 3 degrees (which, by linear distance, is closer to the fovea, in a shorter eye than in a longer eye) increased with eye length, but the trends were not significant. Li, et al. (2010) made similar measures, but closer to the fovea (from 0.10 mm to 0.30 mm eccentricity). They found that linear cone density decreased with eye length, but the trends were not significant at the smallest eccentricities (0.1 and 0.2 mm). When the data were plotted in angular units and angular distance from the fovea, they found that angular cone density trended toward an increase with eye length but none of the trends were significant. A more recent study measured peak cone densities in the fovea as well as axial length for 22 eyes of 22 subjects (Wilk et al., 2017) but they did not plot peak cone density as a function of axial length, as it was not the aim of their study. We plotted the data they provided in their paper and found that the linear cone density at the foveal center dropped significantly with increases in axial length, similar to what was found by Li et al. (2010) and Chui et al. (2008), but the angular cone density had no dependency on eye length. Summary plots from previous literature are shown in Figures 2ab.

**Figure 2.**
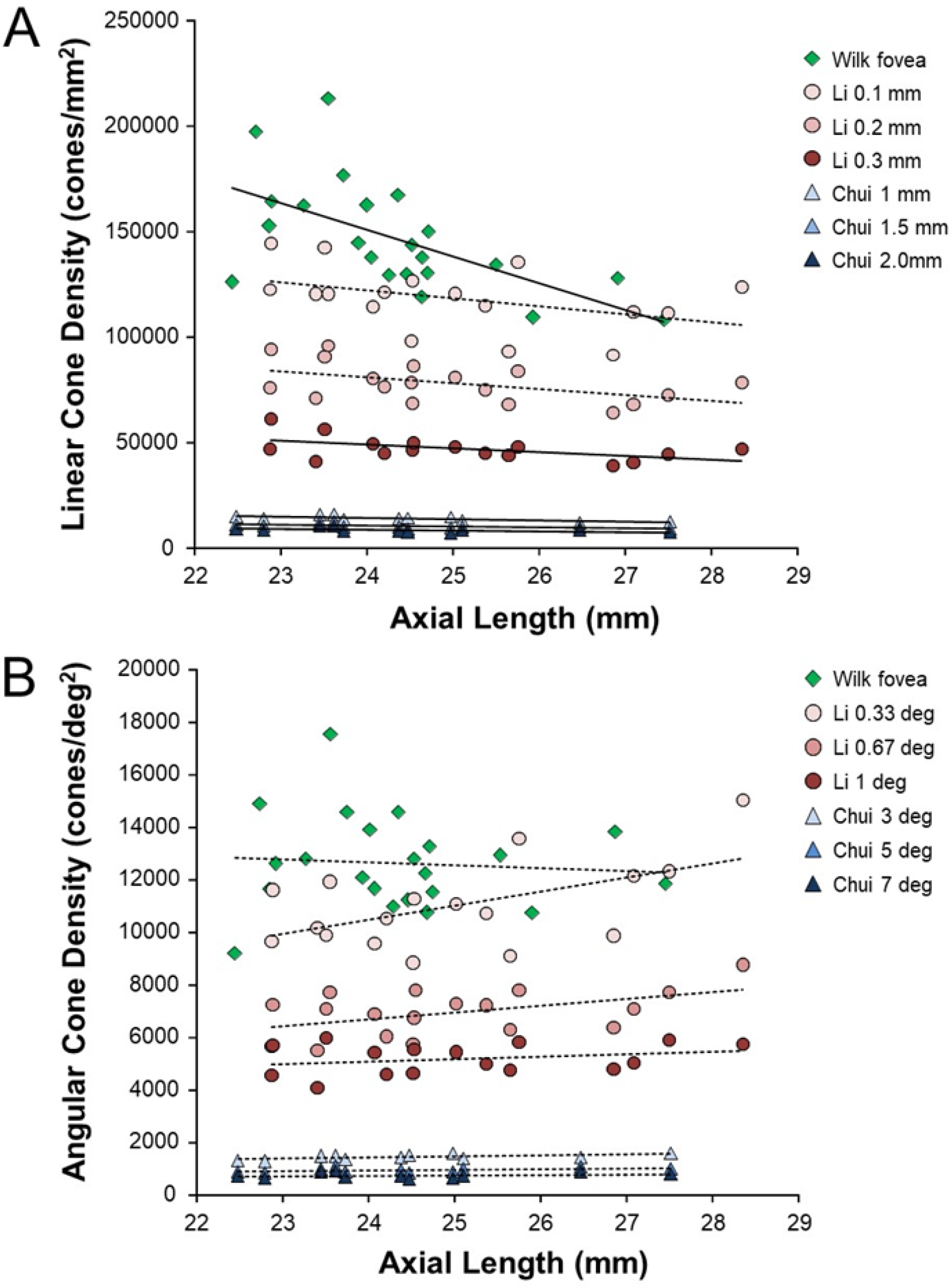
Summary of published data from Li et al. (2010), Chui et al. (2008) and Wilk et al. (2017). In both plots, the linear fits with the solid lines indicate the data that have significant trends. (a) Linear cone density has a decreasing trend with axial length near the fovea. **(b)** Angular cone density (sampling resolution) of the eye generally increases with axial length although none of the data show a significant linear relationship.

Wilk et al. (2017)’s data were consistent with a global expansion model and Li et al. (2010) and Chui et al. (2008)’s data only leaned toward a model that falls between the global expansion and equatorial stretching models. If the trends found by Li et al. (2010) and Chui et al. (2008) near the fovea were to extend to the foveal center, then myopes would have higher foveal photoreceptor sampling resolution with a consequent potential for better performance on visual tasks compared to emmetropes. As such, the simplest explanation for visual deficits in myopes –increased separation between cones caused by retinal stretching – would have to be ruled out.

With the improvements in resolution of adaptive optics ophthalmoscopes, imaging the smallest cones at the foveal center is now possible in many eyes, enabling a definitive analysis of the cone density at the fovea as a function of eye length.

## Results

The experiments were approved by the University of California, Berkeley Committee for the Protection of Human Subjects. All subjects provided informed consent prior to any experimental procedures. Subjects self-reported their eye health so that only healthy individuals with no ocular conditions were included in the study. All eyes were dilated and cyclopleged with 1% Tropicamide and 2.5% Phenylephrine before imaging. We report data from 28 eyes of 16 subjects with a wide range of refractive error and axial length. Age, sex and ethnicity are listed on Table 2.

**Table 2.**
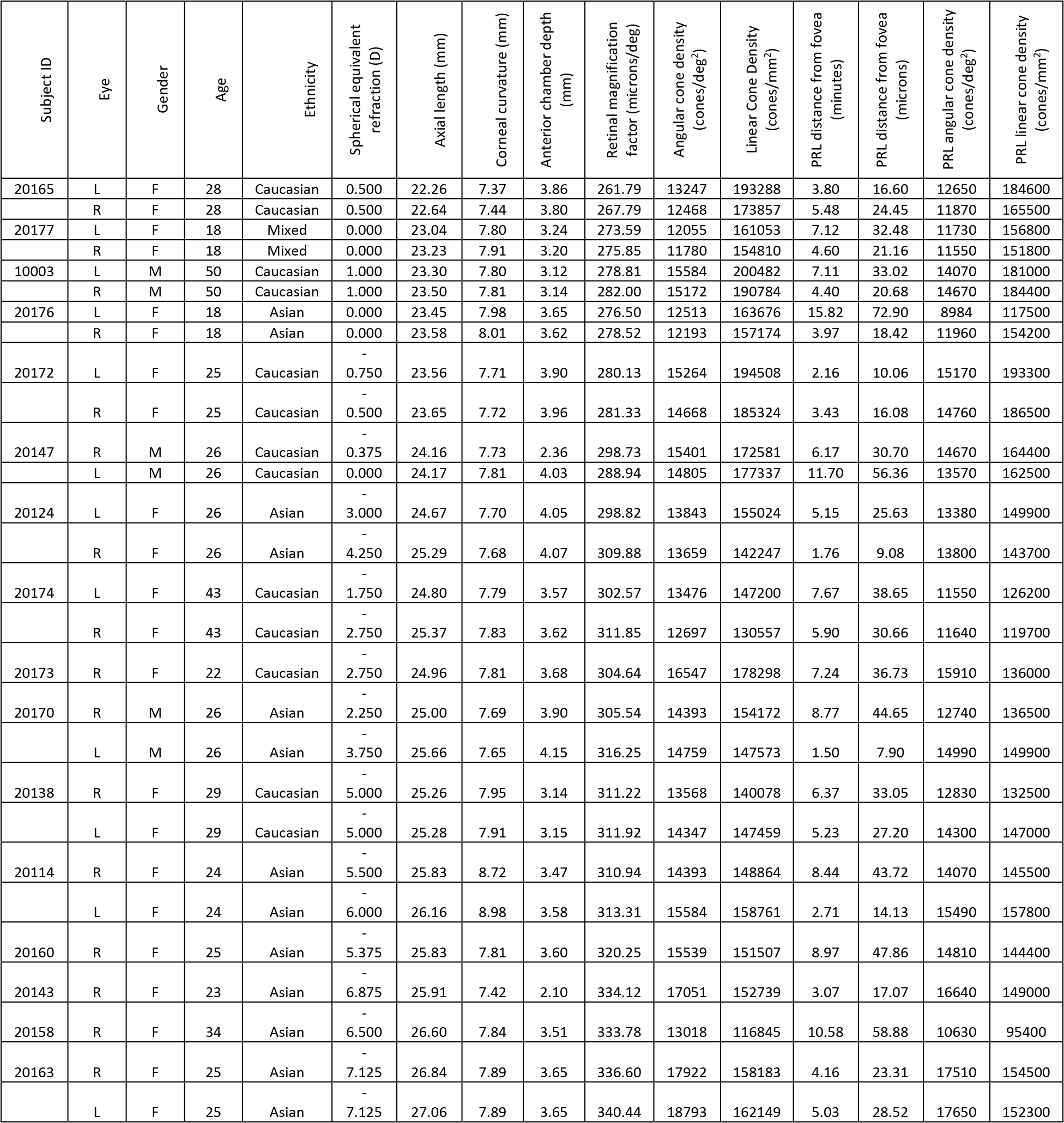
Each subject’s refractive error was self-reported at the time of the study. Axial Length, corneal curvature and anterior chamber depth were measure by IOL Master, and retinal magnification factor (microns/deg) was calculated from biometry data.

### Biometry Data

All the biometric measures used to convert angular dimensions to linear retinal dimensions are listed on Table 2. The strong correlation between refractive error and eye length (P < 0.0001) indicates that the myopia was predominantly as a result of axial length.

### Imaging Data

Images of the foveal region, the preferred retinal locus for fixation (PRL) and the fixation stability were recorded with an adaptive optics scanning laser ophthalmoscope (see **Methods and Materials**). The image of one subject (10003L) is shown in Figure 3a. All the cones were resolved with our imaging system. The scatter plot indicates the scatter plot of fixation over the course of a 10-sec video. Figure 3b shows the same image with all cones labeled and a color-coded overlay indicating the density. 16,184 labeled cones are shown on the figure. The point of maximum density is indicated by the blue cross and the average location of the PRL is indicated by the yellow cross (mean of the scatter plot locations in Figure 3a). This eye has a peak linear density of 200,482 cones/mm^2^, and a peak angular density of 15,584 cones/deg^2^. Cone density plots in linear and angular units for all eyes are shown on supplemental figures 1 and 2. Original images and a list of the cone locations for each can be downloaded from the Resources section of the Roordalab website (roorda.vision.berkeley.edu).

**Figure 3.**
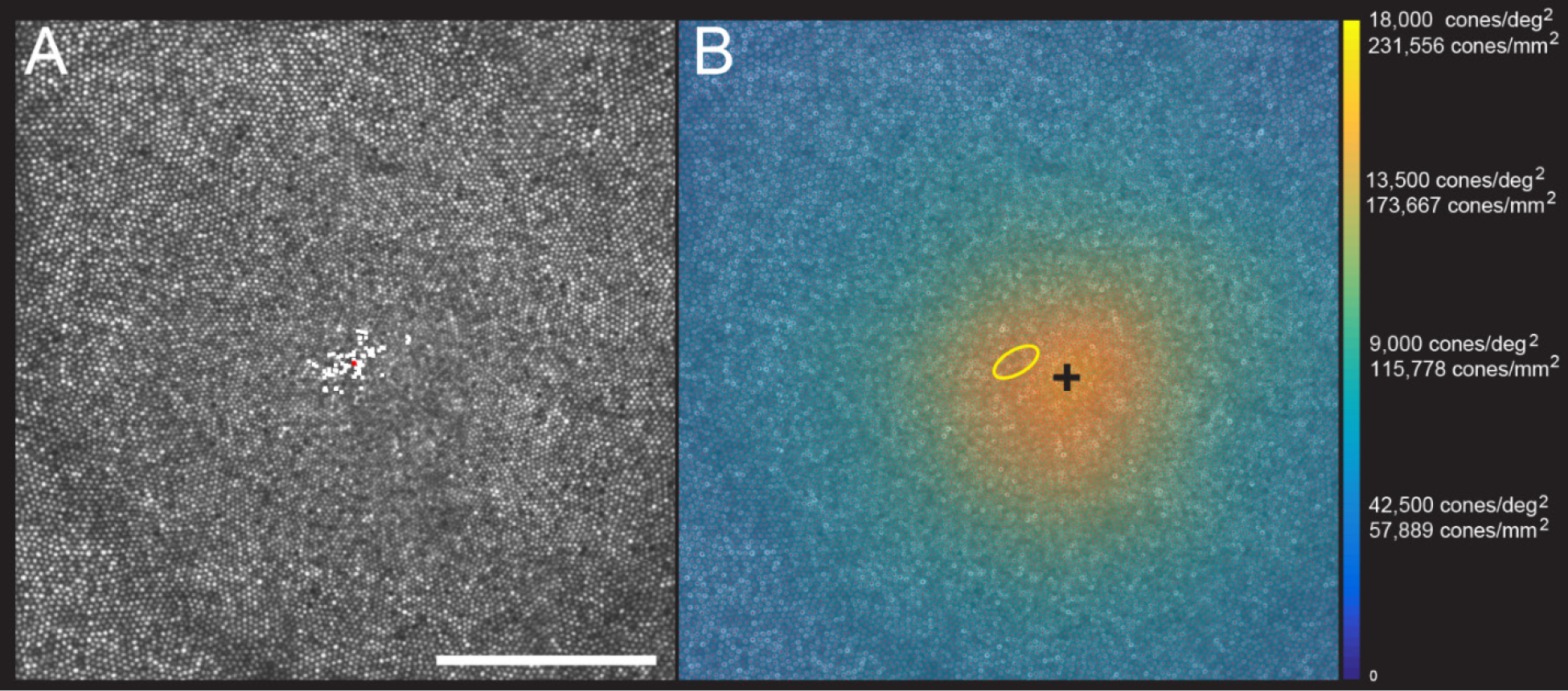
**(a)** AOSLO image of the fovea one subject (10003L). Only the central 1.5 degrees are shown here (810 X 810 pixels), which contains 16,184 cones. The white dots are a scatter plot showing the PRL, or position of the fixated stimulus over the course of a 10-second video. The red dot is the centroid of the scatter plot. **(b)** Same image with a color overlay indicating the density. Linear and angular cone densities are indicated on the right colorbar. Peak cones densities in this eye are 200,482 cones/mm^2^ and 15,584 cones/deg^2^. The yellow ellipse is the best fitting ellipse containing ~ 68% of the points in the scatterplot and indicates the PRL. The black cross indicates the position of peak cone density. Scale bar is 0.5 degrees, which in this eye corresponds to 139.4 microns.

Figure 4 shows the linear cone density as a function of linear eccentricity, where the average linear cone density was computed in 25-micron wide annuli centered around the point of peak density.

**Figure 4.**
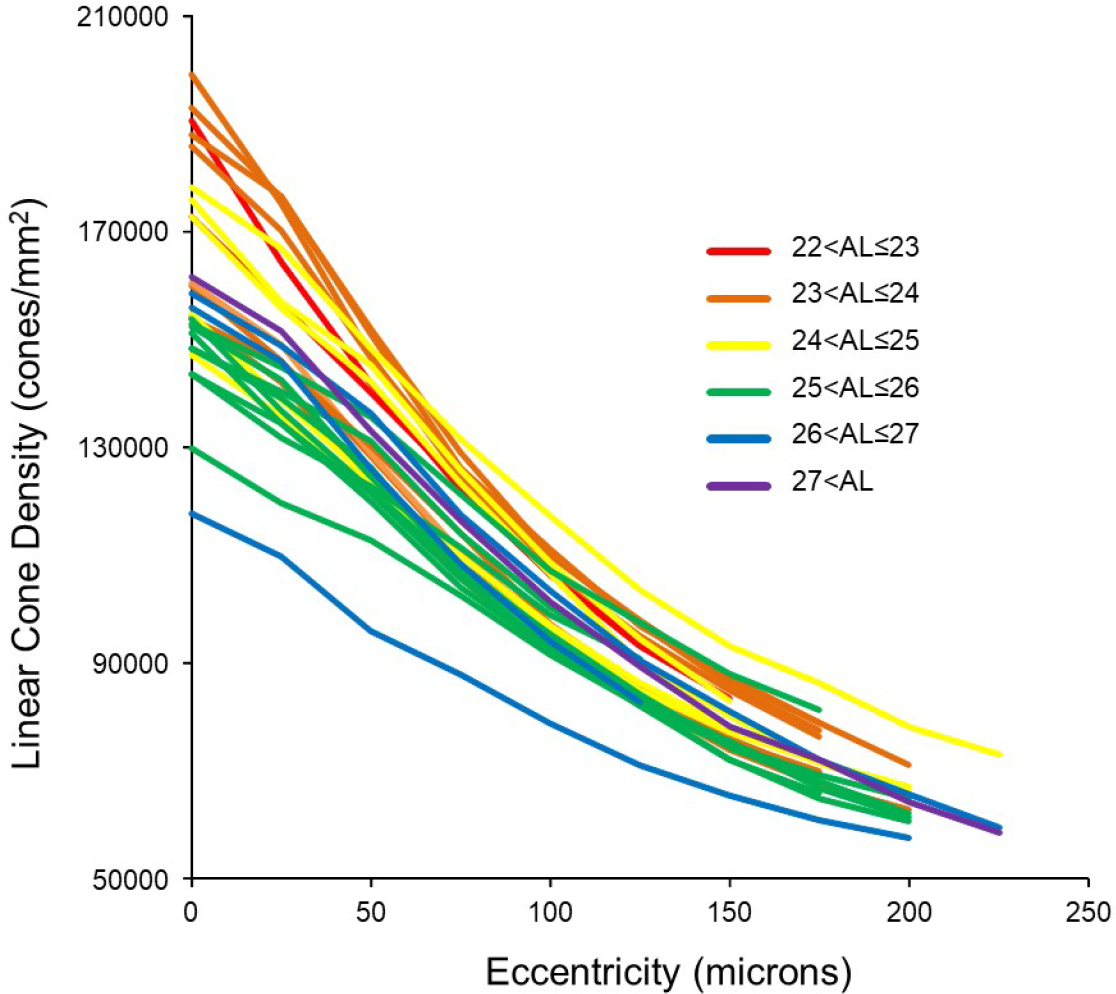
Cone density as a function of eccentricity for all eyes. The axial length ranges of the subjects are color coded, with warmer colors for shorter eyes and cooler colors for longer eyes. In this plot, it is apparent that shorter eyes generally have higher peak cone densities.

In order to show the trends of density with axial length Figure 5a&b plot linear and angular cone density as a function of axial length where the colors indicate different eccentricity - red to purple indicate distance from the from fovea towards more parafoveal locations. Figure 5a reveals that peak linear density decreases significantly with axial length and the trend persists and remains significant from the fovea out to 100 microns eccentricity. Axial length accounts for 38% of the variance in the changes in linear cone density. Figure 5b shows the opposite trends when plotted in angular units. Peak angular density increases significantly with axial length and the trend persists and remains significant out to 40 arcminutes eccentricity. Axial length accounts for 32% of the variance in the changes in angular cone density. The plots clearly indicate that although stretching does occur (Figure 5a) it is not a simple global expansion and longer eyes have higher sampling density. The trends hold at and around fovea with statistical significance.

**Figure 5.**
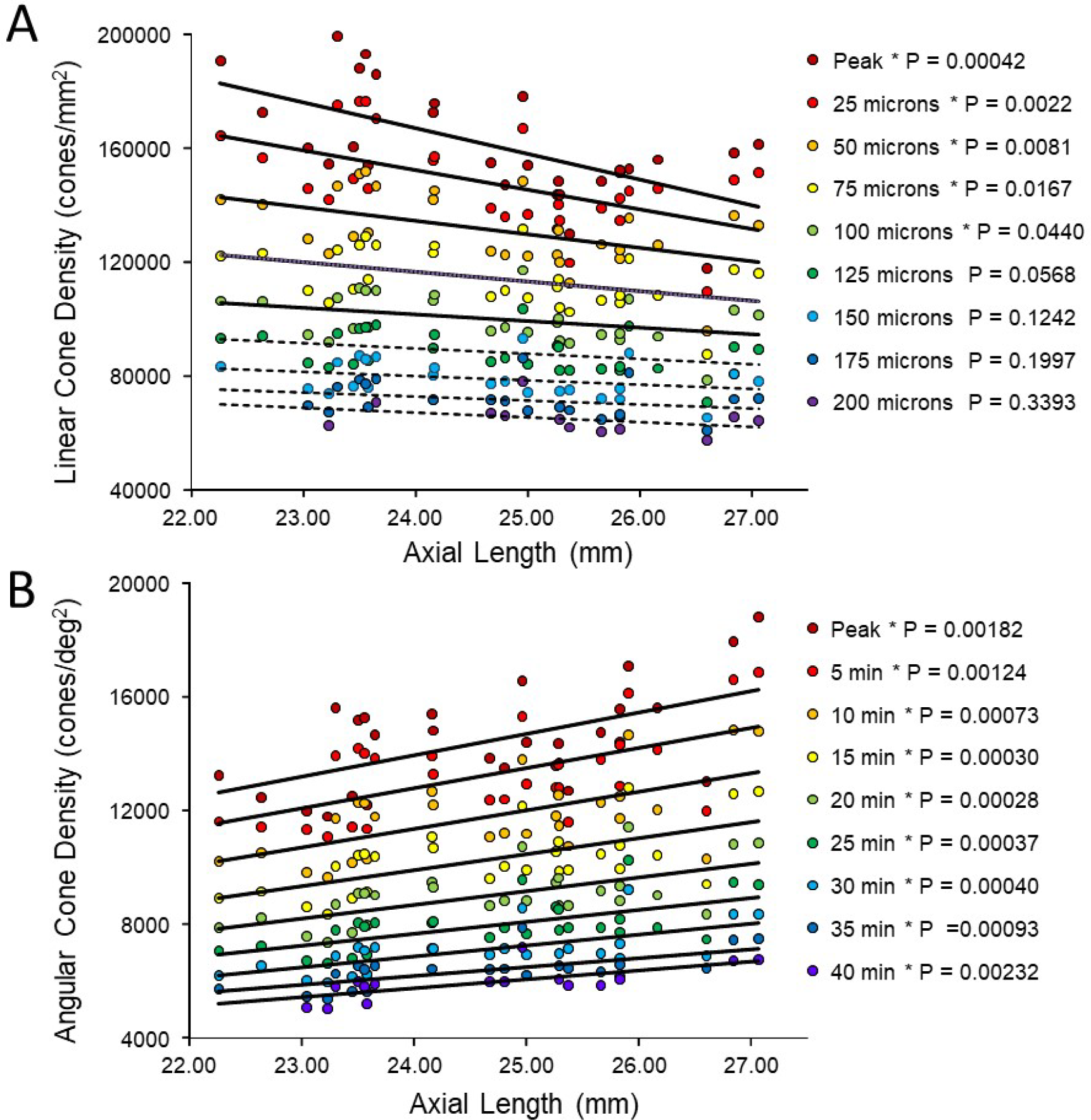
**(a)** Linear cone densities as a function of axial length. Longer eyes have lower linear cone density than shorter eyes. The trend remains significant out to 100 microns eccentricity and **(b)** Angular cone densities as a function of axial length. The peak angular cone density increases significantly with increasing axial length and this trend remains significant out to 40 arcminutes eccentricity. Relationships with P-values <0.05 are labelled with asterisks and trendlines are shown as solid lines. Relationships with P-values ≥0.05 have dashed trendlines.

A more relevant measure of the impact of eye length on vision is how the angular cone density changes at the PRL, which is often displaced from the location of peak cone density (Li et al., 2010; Putnam et al., 2005; Wilk et al., 2017). If, for example, longer eyes had more displaced PRLs then that could diminish, or even reverse, the trend of increased angular density with eye length reported in Figure 5b. We found that the average displacement between PRL and maximum cone density was 5.82 arcminutes and 28.94 microns. There was no significant linear relationship found between PRL displacement in either angular or linear units vs. axial length. Therefore, the PRL was not more displaced in myopes than in emmetropes from the point of peak cone density. Plots of the cone density at the PRL with axial length show the same trend at the PRL as at the point of maximum cone density (Figure 6 a&b).

**Figure 6.**
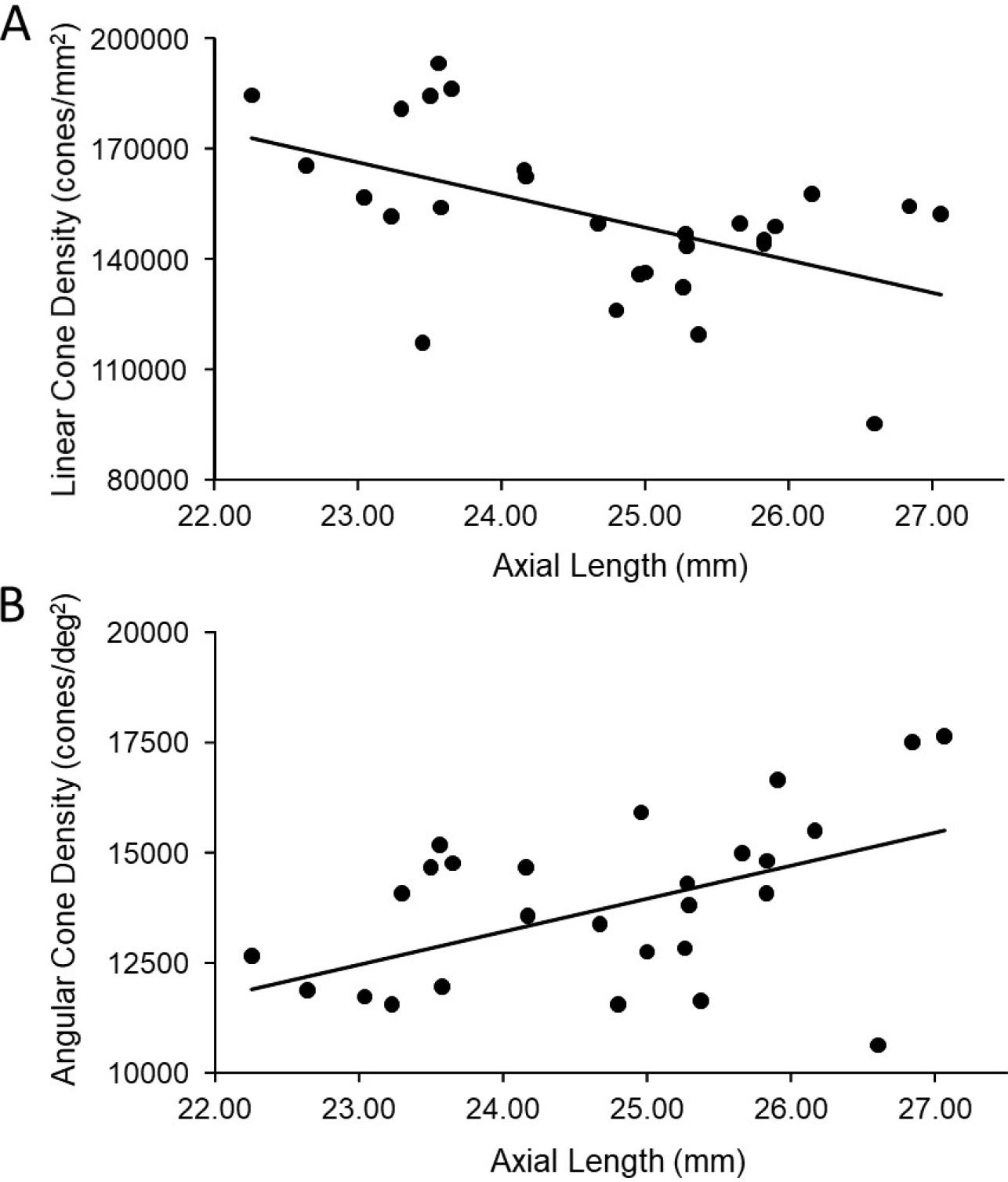
**ab.** The relationship between cone density and axial length shows the same pattern at the PRL as for the peak cone density. The slopes in both (a) and (b) are significant (P=0.00975 & P=0.00432 respectively) and axial length accounts for 23% and 27% of the variance in linear and angular cone density, respectively.

Finally, we explored whether fixational eye movements might have a dependency on axial length. Fixation stability around the PRL had an average standard deviation of 3.94 arcminutes and 19.84 microns. The average area of the best fitting ellipse containing ~ 68% of the points in the scatterplot (defined as the bivariate contour ellipse area, or BCEA) was 50.7 square arcminutes and 1303 square microns. The plot of BCEA in square microns v.s. axial length v.s. showed a trend that approached significance (P=0.0596) (Figure 7a), but when we plotted BCEA in square arcminutes v.s. axial length, the trend was no longer apparent (P=0.364) (Figure 7b). In other words, if there is any increase in fixational eye movements in microns, it is just a symptom of having a longer eye.

**Figure 7.**
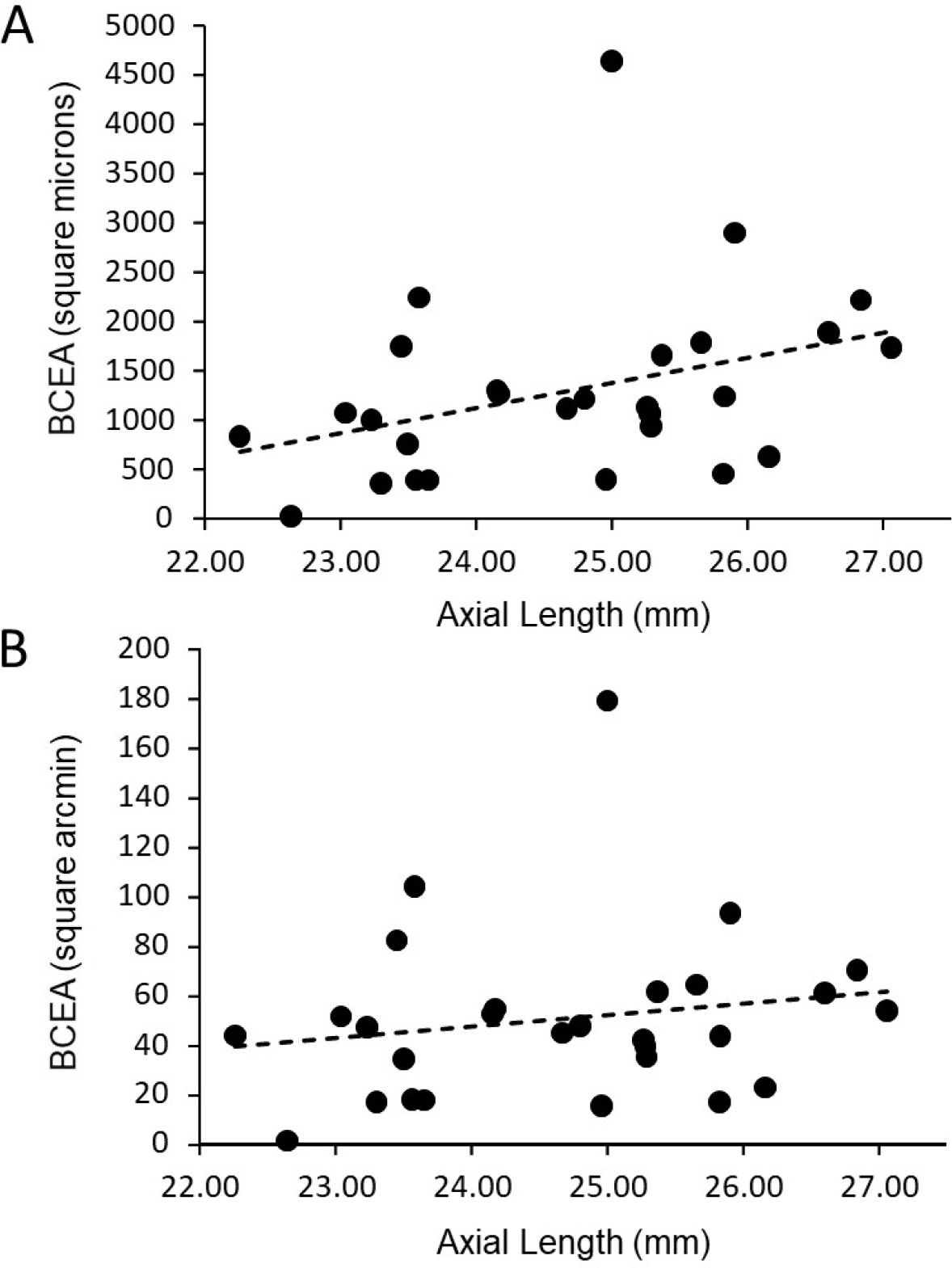
**(a)** The plot of BCEA in linear units (square microns) v.s. axial length shows a trend that approaches significance (P=0.0596) **(b)** There is no significant relationship between BCEA in angular units (square arcminutes) and axial length (P=0.364).

## Discussion

In this paper we measure the cone density at and near the foveal center and investigate how it changes as a function of axial length. This is the first comprehensive study of cones in living eyes at the foveal center, the area solely responsible for a human’s fine spatial vision. Our results show that although some expansion does occur (linear cone density decreases with axial length) the angular sampling resolution actually increases, on average, with axial length. Prior to this study, the relationships between cone density and axial length were only made outside of the fovea, the closest being 0.1 mm, or 0.3 degrees (Li et al., 2010). Although an eccentricity of 0.3 deg might seem close, it is noted that the cone density drops precipitously just outside of the location of peak density (Curcio, Sloan, Kalina, & Hendrickson, 1990) as does human vision (Poletti, Listorti, & Rucci, 2013)(Rossi & Roorda, 2010b). There are other factors that govern peak cone density, however; eye length accounts for anywhere between 27% and 38% of the variance in cone density.

Our finding that the slopes of cone density vs. axial length are in opposite directions when plotted in linear (negative slope) and angular (positive slope) units, supports an eye growth model that lies between the global expansion model and an equatorial stretching model. Previous studies from our lab (Li et al., 2010) and also from Chui et al. (2008) leaned in the same direction. None of the cone density studies provide insight into the reasons why the photoreceptor density would behave this way with eye growth, but the results do align with other observations reported in the literature. Specifically, Atchison et al. (2004) used magnetic resonance imaging and found that eyeball dimensions in axial myopes are variable but are generally larger in all directions with a weak tendency to be preferentially greater in the axial direction. Their reported eye growth patterns lie between that illustrated for the global expansion and equatorial stretching models in figure 1.

Our results differ from Wilk et al. (2017) whose data support a global expansion model (i.e. there is no detectable change in angular cone density with axial length; figure 2b). But it is important to point out that their study did not set out to address the same question and the number of subjects with long axial lengths was disproportionately low.

Our results also differ from Troilo (1998) who studied retinal cell topography in a marmoset animal myopia model. Higher cone packing densities were observed in the experimentally enlarged eyes compared to normal eyes in the fovea. Their result followed the overdevelopment model, which is the reason why we included it as one of the possible outcomes of our study. In fact, the overdevelopment model is an extension of Springer’s model of development (Springer & Hendrickson, 2004), which offers a biomechanical explanation for how cone packing increases at the foveal center in a developing eye. While our data do not support the overdevelopment model, it does not preclude the existence of biomechanical factors working in opposition to simple global expansion.

The fact that angular cone density (visual sampling resolution) increases with eye length (myopia), at the peak density and at the PRL, means that poorer performance by myopes on resolution tasks cannot be explained by a decrease in photoreceptor sampling. The deficit musts arise at a post-receptoral level.

Low-level causes for myopic visual deficits might arise from differences in the connectivity between cones and ganglion cells. Atchison et al. (2006) suggested that abnormal eye growth may be associated with a loss of ganglion cells. Alternately, if ganglion cells pool signals from multiple cones, then they will impose the retinal sampling limit and reduce certain aspects of visual performance (acuity, for example). Recent electron microscopy studies of a human fovea have revealed extensive convergence and divergence connections between photoreceptors and ganglion cells, albeit in an eye from an individual who was born prematurely (Dacey, 2018). These discoveries challenge our current understanding of neural connectivity in the foveal center and force us to consider the possibility of interindividual differences in foveal cone wiring. More experiments are necessary to explore these ideas.

To explain why low myopes did not perform as well on an acuity task as emmetropes, even after correction or bypassing of high order aberrations, Rossi et al. (2007) and Coletta & Watson (2006) both raised the possibility that myopes might have become desensitized to high frequency information (low level myopic amblyopia) as a result of having less exposure to a high contrast visual environment. In this case, it might be possible to train myopes to take advantage of their higher sampling resolution, but one myope in a follow up study by Rossi & Roorda (2010a) never reached the acuity levels of emmetropes in the same study.

### Comparisons with Previous Studies

#### Peak cone densities

Curcio et al. 1990 measured spatial density of cones and rods in eight explanted whole-mounted human retinas. They found a large range of peak foveal cone densities with an average of 199,000 cones/mm^2^. When we averaged the peak cone density over a circular aperture of 7.5 arcminutes which was similar to the 29 x 45 micron window that Curcio et al. (1990) used to compute density, we measured peak linear cone densities ranging from 123,611 to 214,895 with an average of 168,047 cones/mm^2^. Zhang et al. (2015) reported an average peak density of 168,162 cones/mm^2^ in 40 eyes although they used a much smaller 5 x 5 micron sampling window to measure the peak. Wilk et al. (2017) reported an average peak density of 145,900 cones/mm^2^ in 22 eyes using a 37 x 37 micron sampling window and Li et al. (2010) reported an average peak density of 150,412 cones/mm^2^ in 4 eyes over a sampling window encompassing 150 cones (approximately 37 micron diameter at the foveal center). All reports of cone densities from adaptive optics studies in living eyes are lower than reports from histology. Two possible reasons for this are (i) the excised tissue in Curcio et al. (1990) underwent more shrinkage than estimated or (ii) the adaptive optics reports are subject to selection bias, where individuals with the highest angular cone densities might have been excluded because the image were less well resolved rendering the cones images too difficult to label with confidence. In our study, we attempted to image 73 eyes from 46 subjects and only succeeded in resolving cones across a sufficiently large region at and around the fovea in 28 of them. The reason the images from 45 eyes were not analyzed was due to poor or inconsistent image quality arising from a number of factors: Images from 4 eyes (3 subjects) were not analyzed because their refractive errors were too high (all above –8D) and we ran into the limits of the deformable mirror’s dynamic range. Images from 18 eyes (13 subjects) that were taken early on in the study were not analyzed because the optics of AOSLO were not tuned well enough to resolve foveal cones. Images from 4 eyes (2 subjects) were not analyzed because of uncorrectable image degradation caused by keratoconus and corneal scarring. Images from 2 eyes (1 subject) were not analyzed because of excessive aberrations caused by an orthokeratology refractive correction. The cause of poor or inconsistent image quality among the remaining 17 eyes were varied, including ocular surface dryness, excessive eye motion and small pupils. The average refractive error among these remaining 17 eyes was about the same as the successful eyes.

#### Anisotropic density distribution

Like Curcio et al. (1990) and Zhang et al. (2015) we found steeper drops in cone density in the superior and inferior directions compared to the nasal and temporal directions. Plots of density along the two cardinal directions are shown on Supplemental Figure 3.

#### PRL displacements

The distance of the PRL from the foveal center for our study (mean 29 microns; range 8 – 73; n = 28) roughly agrees with those of Wilk et al. (2017) (mean 63 microns; range 20 – 263; n = 22), Li et al. (2010) (mean 34 microns; range 3 – 92; n = 18) and Putnam et al. (2005) (mean 17; range 11 – 23; n = 5). The differences in cone density between the peak and the PRL were small and the trends (Figures 5 and 6) persisted at both locations.

#### Spatial vision estimates

The cone array imposes the first retinal sampling limit to human spatial vision (MacLeod, Williams, & Makous, 1992; Williams, 1985) and the photoreceptor row-to-row spacing (assuming an hexagonal packing structure) imposes the maximum frequencies that can be relayed to later stages without aliasing. We can compute the sampling limit and estimate the cone center-to-center spacing using the following formulas:

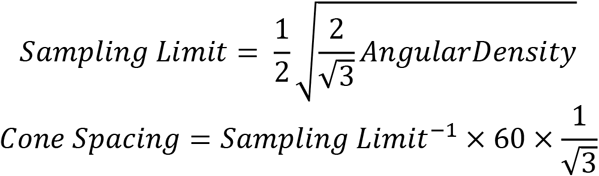

For the densities reported here, the potential spatial frequency resolution limits range from 58.3 to 73.6 cyc/deg (average: 64.5 cyc/deg) at the peak density and 50.9 to 71.4 cyc/deg (average: 62.7 cyc/deg) at the PRL. These correspond to potential acuities ranging from 20/11.8 to 20/8.2 (based on the primary spatial frequency of the three bars of a Snellen E). The cone frequency cut-offs are higher than almost all the interferometric acuity limits reported by Coletta & Watson (2006), even for the emmetropic subjects. The acuities are, however, in the range of those measured from emmetropic subjects after adaptive optics correction by Rossi et al. (2007). The cone center-to-center spacing ranges from 0.59 to 0.47 arcminutes at the peak density and from 0.60 to 0.49 arcminutes at the PRL. A direct comparison of foveal structure and function for each of our subjects was not the scope of this study but will be the topic of future investigation.

Measuring structure and function of cone photoreceptors at the foveal center – the most important region of the human retina – has been one of the more challenging endeavors in vision science. Fortunately, the latest generation of adaptive optics ophthalmoscopes are making it easier and are facilitating new discoveries within this retinal region. The pattern of how cone density changes with eye growth lands somewhere between the global expansion and equatorial stretching models. The cone mosaic in longer eyes is expanded at the fovea, but not in proportion to eye length. Despite retinal stretching, myopes generally have a higher angular sampling density in and around the fovea compared to emmetropes. Reports of reduced best-corrected central visual acuity in myopes compared to emmetropes cannot be explained by decreased photoreceptor density caused by retinal stretching during myopic progression.

## Materials and Methods

### Foveal Imaging

We used our latest generation adaptive optics scanning laser ophthalmoscope (AOSLO) for foveal imaging. The system used a mirror-based, out-of-plane optical design (Dubra et al., 2011), and employed a deformable mirror with a continuous membrane surface and shaped with 97 actuators (DM97, ALPAO, Montbonnot-Saint-Martin, France). The system scans multiple wavelengths simultaneously. Each wavelength was drawn from the same broadband supercontinuum source (SuperK EXTREME, NKT Photonics, Birkerod, Denmark) using a custom-built fiber coupler. Wave aberrations were measured with a custom-built Shack Hartmann wavefront sensor using the 940 nm channel. Images were recorded using the 680 nm channel. 512 x 512 pixel videos were recorded over a 0.9 x 0.9 degree square field for an average sampling resolution of 9.48 pixels per arcminute. Eye alignment and head stabilization was achieved by using either a bite bar or a chin rest with temple pads. At least one 10-second video was recorded at the fovea and at 8 more locations where the subjects were instructed to fixate on the corners and sides of the raster, to image an entire foveal region spanning about 1.8 X 1.8 degrees. In order to ensure the best possible focus of the foveal cones, multiple videos were taken over a range of 0.05 D defocus steps to find the sharpest foveal cones. Focus steps were generated by adding a focus shape onto the deformable mirror. Online stabilization and registration algorithms were used to facilitate rapid feedback on the image quality.

### Locating the Preferred Retinal Locus of Fixation (PRL)

Steady fixation was achieved at the fovea center by having the subjects fixate on a dark, circular, blinking dot with a diameter of 3.16 arcminutes (30 pixels) in the center of the AOSLO scanning raster. The fixation target was generated by modulating the same 680 nm scanning beam used for imaging and, as such, the target’s location was encoded directly into each frame of the video (Poonja, Patel, Henry, & Roorda, 2005). A scatter plot of the positions of the blinking dot relative to the retina was generated and was fit with a bivariate ellipse using free online Matlab scripts downloaded from http://www.visiondummy.com/wp-content/uploads/2014/04/error_ellipse.m. The bivariate contour ellipse area (BCEA), which is the area of the best-fitting ellipse encompassing 68% of the points in the scatterplot (Castet & Crossland, 2012) was used to quantify the fixation stability (figure 7) and the exact location of the PRL within the imaged cone mosaic (Table 2, Figure 3, Supplemental figures 1 and 2).

### Image Processing and Analysis

High quality images were generated from the recorded videos offline using custom software (Matlab, The MathWorks, Inc., Natick, MA) to measure and correct for distortions caused by eye movements (Stevenson & Roorda, 2005). Poor-quality frames were manually excluded and registered frames were averaged into a single high signal-to-noise image. The processed images were stitched together (Photoshop; Adobe Systems, Inc., Mountain View, CA) to create an approximately 1.8-degree montage of the foveal cone mosaic.

We used custom software to identify and label individual cones in the AO retinal images. The program allows the user to select a region of interest and manually add and delete cone labels. A combination of both manual and automated methods (Li & Roorda, 2007) were used to identify cone locations as the current version of the program does not adequately recognize cones in the foveal center where they are dim and smaller (Li et al., 2010). All the cone coordinates were selected and reviewed by two of the authors. In some cases cones were too dim to be seen but there was only a gap in the mosaic (Bruce et al., 2015). If a space that might have been occupied by a cone was dim or dark, we would assume it was a cone and mark its location. We rationalize this for two reasons: First, if there is a gap in the mosaic, then it is likely that a cell is occupying that space, otherwise the adjacent cells would migrate to fill it in (Scoles et al., 2014). Second, in our experience and of others (Pallikaris, Williams, & Hofer, 2003), cones that appear dark in one visit, can often appear bright in the next. In other cases (uncommon) the contrast was low in some regions or there were interference artifacts in the images (Meadway & Sincich, 2018; Putnam, Hammer, Zhang, Merino, & Roorda, 2010), making the cone locations slightly ambiguous. In these instances, we made manual cone selections based on the assumption that the cones were all similar in size and close-packed into a nearly hexagonal array (Curcio et.al., 1990).

Continuous density maps were generated by computing cone density within a circle of 10 arcminutes in diameter around every pixel location across the image. We kept the area large enough to generate smooth maps, but small enough to resolve local changes. Changes in density with eccentricity were generated by computing the density in 5 arcminute annuli surrounding the point of peak cone density. For linear density measures we used annuli with 25 micron widths.

### Retinal Magnification Factor Calculation

The exact angular dimensions of the AOSLO images were computed by imaging a calibrated model eye in the AOSLO system, but the conversion to linear dimensions on the retinal image requires additional measurements, since the dimensions of each eye governs the actual size of the image on its retina. The conversion from visual angle to retinal distance requires a measurement of the axial length of the eye and an estimation of the location of the secondary nodal point. We used a four-surface schematic eye model, originally proposed by Li et al., 2010 to estimate the location of the secondary nodal point. The corneal first surface radius of curvature, the anterior chamber depth and the axial length were for measured for each subject with an IOL Master (Zeiss Meditec, Dublin, CA). The radius of the curvature of the back surface of the cornea was computed as 88.31% of the front surface (Bennett, Rudnicka, & Edgar, 1994). The indices of refraction of the media and the radii of curvature of the front and back lens surface were taken from the Gullstrand schematic eye (Vojnikovic & Tamajo, 2013). Once determined, retinal image size is related to visual angle by the equation:

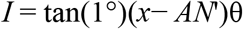

Where *I* is retinal image size, *x* is axial length, *AN*’ is the distance from the corneal apex to the eye’s second nodal point, and θ is the visual angle. As can be seen in Table 2, myopic eyes, which generally have longer focal lengths, have proportionally larger retinal images.

### Statistical Analysis

Given the trends of increased angular density as a function of axial length that Li et al (2011) observed at the location closest to the fovea (slope = 531 cones/deg^2^ for each mm increase in axial length; standard deviation of the regression errors = 1377 cones/deg^2^), we estimated that data from approximately 32 eyes, evenly distributed across a range of axial lengths would be sufficient to show if there was a true effect at the fovea. The targeted number was computed using methods outlined by Dupont & Plummer (1998) implemented using free online software (Power and Sample Size Program Version 3.0, January 2009, downloaded from http://biostat.mc.vanderbilt.edu/wiki/Main/PowerSampleSize) with type 1 error probability of 0.05 and a power of 0.95.

All data collected in this study were analyzed using simple linear regression models in Excel. P-values for all linear regressions are reported and linear trendlines with P-values less than 0.05 are plotted as solid lines and P-values greater than 0.05 as dashed lines.

---

**Supplemental Figure 1:**
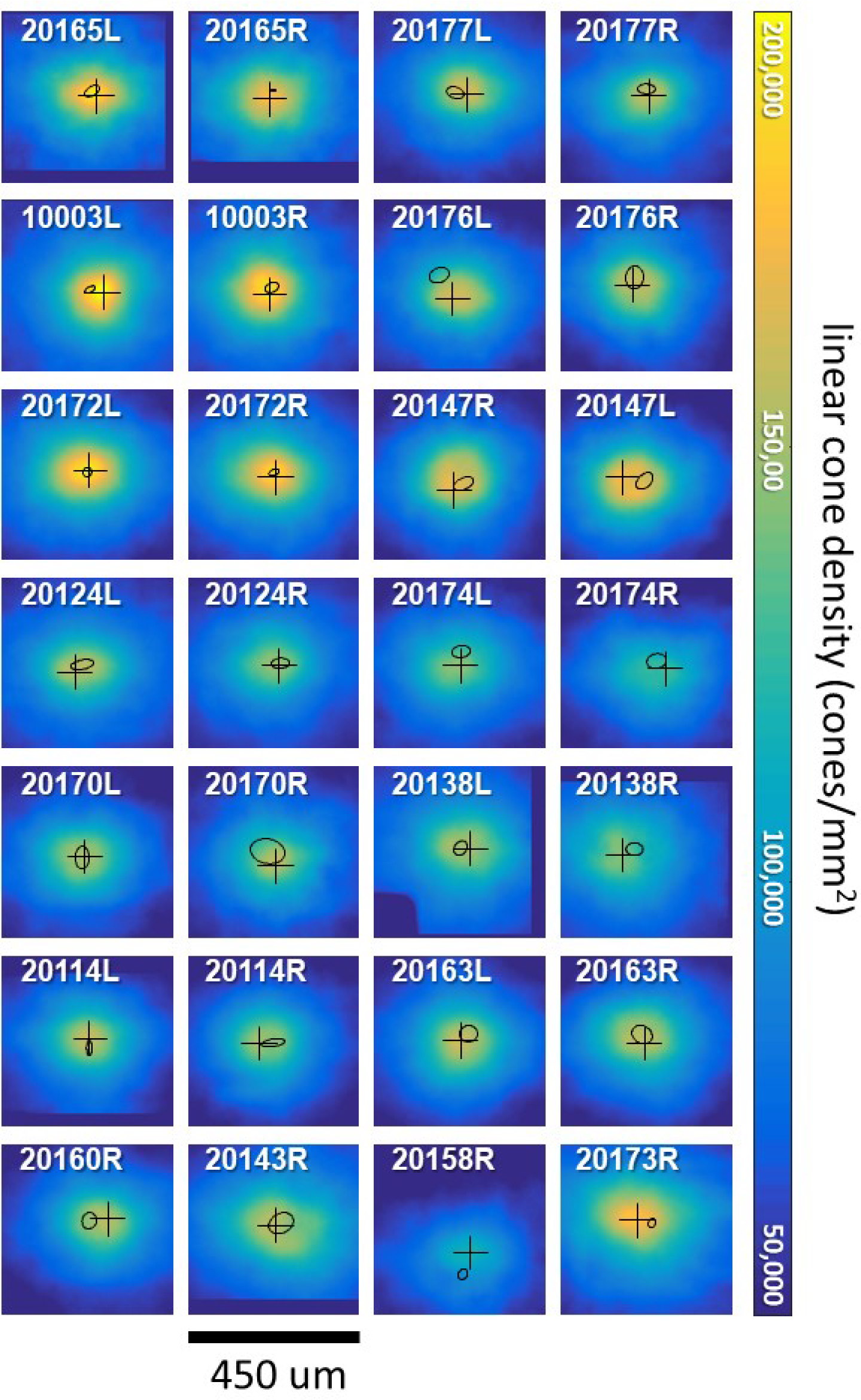
Linear cone density (cones/mm^2^) plots over the central 450 microns for all 28 eyes. The black cross indicates the point of maximum cone density. The black ellipse is the best fitting ellipse about the fixation scatterplot indicating the PRL. Dark blue regions indicate where no cone density estimates were made.

**Supplemental Figure 2:**
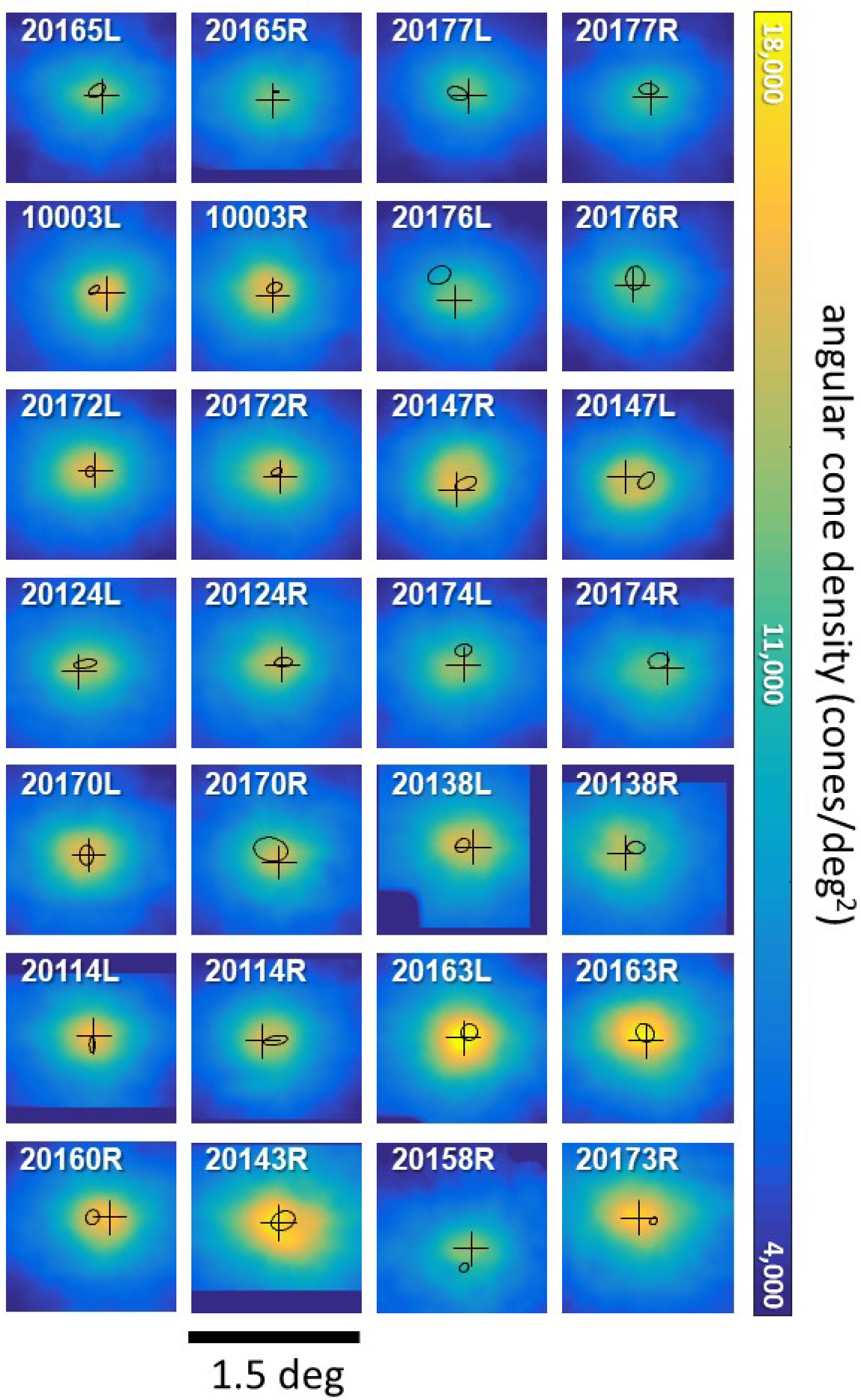
Angular cone density (cones/deg^2^) plots over the central 1.5 degrees for all 28 eyes. The black cross indicates the point of maximum cone density. The black ellipse is the best fitting ellipse about the fixation scatterplot indicating the PRL. Dark blue regions indicate where no cone density estimates were made.

**Supplemental Figure 3:**
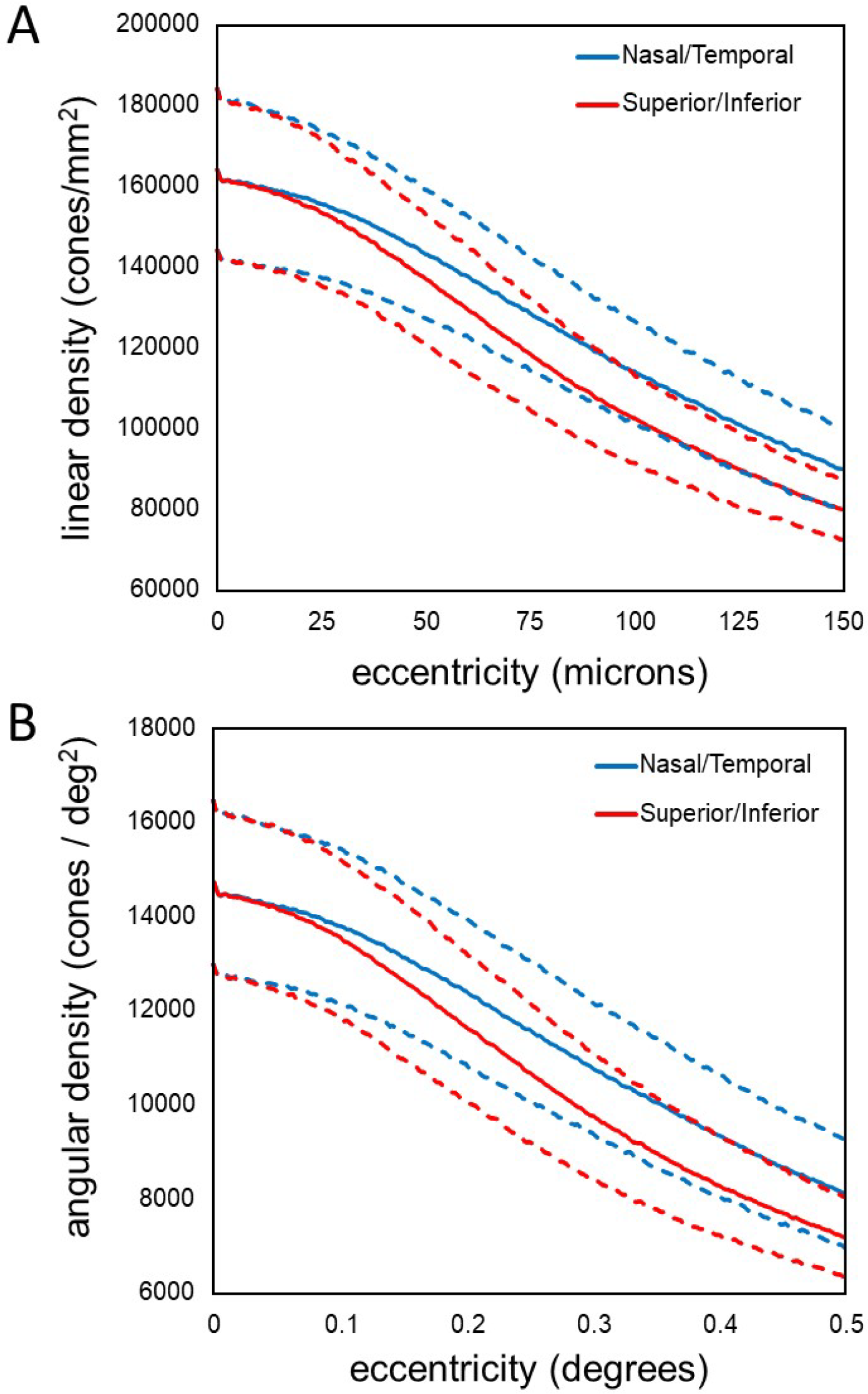
Plots of density as a function of eccentricity in the vertical and horizontal directions. (A) linear cone density (B) angular cone density. The dashed lines represent +/− 1 standard deviation from the mean.

